# Identification of key genes in SARS-CoV-2 patients on bioinformatics analysis

**DOI:** 10.1101/2020.08.09.243444

**Authors:** Hanming Gu, Gongsheng Yuan

## Abstract

The COVID-19 pandemic has infected millions of people and overwhelmed many health systems globally. Our study is to identify differentially expressed genes (DEGs) and associated biological processes of COVID-19 using a bioinformatics approach to elucidate their potential pathogenesis. The gene expression profiles of the GSE152075 datasets were originally produced by using the high-throughput Illumina NextSeq 500. Gene ontology (GO) and Kyoto Encyclopedia of Genes and Genomes pathway (KEGG) enrichment analyses were performed to identify functional categories and biochemical pathways. GO and KEGG results suggested that several biological pathways such as “Fatty acid metabolism” and “Cilium morphogenesis” are mostly involved in the development of COVID-19. Moreover, several genes are critical for virus invasion and adhesion including FLOC, DYNLL1, FBXL3, and FBXW11 and show significant differences in COVID-19 patients. Thus, our study provides further insights into the underlying pathogenesis of COVID-19.

## Introduction

The COVID-19 is a major threat worldwide, which lacks effective medication^1^. Moreover, the understanding of mechanisms of anti-virus immunity is limited^2^. COVID-19 genome sequence homology with SARS-CoV is 77% and has the potential to become a seasonal disease^3^. COVID-19 is a troubling disease with the feature of high infection rate, long incubation period, and atypical symptoms^4^. The world is not prepared for the pandemic and more infections have appeared in the second phase of the outbreak^5,6^. Thus, the rapid development and administration of vaccines and drugs are of great importance to quell this pandemic.

The COVID-19 is an RNA virus with spike-like glycoproteins^7^. The major structural proteins of coronaviruses are spike protein, envelope protein, membrane protein, and nucleocapsid protein^8^. Based on these structures, DNA/RNA vaccines in situ inside the patient should be considered^9^. Modern virology and antiviral drug discovery are expected to be principal by analyzing genomics methods^10^. The future challenge is to determine which gene products are essential for virus survival or disease progression^11^. Thus, exploring the unique characteristics belonging to COVID-19 is principal in developing therapies to improve patient outcomes.

A great number of strategies have been taken to explore the molecular characteristics of COVID-19^12^. High-throughput microarray methodologies and advanced drug development including remdesivir and acyclovir have drawn extensive attentions^13^. Moreover, multiple gene expression profiling studies on COVID-19 have been performed by using microarray technology^14^. However, due to the shortcomings of microarray such as limited sample size, deficiency of information, and measurement issue, the understanding of key pathogenesis of COVID-19 becomes a major challenge^15^.

In this study, we analyzed the GEO data (GSE152075), provided by Lieberman N and Greninger A (University of Washington), from the Gene Expression Omnibus database (http://www.ncbi.nlm.nih.gov/geo/) to identify DEGs and the relevant biological process of COVID-19 utilizing comprehensive bioinformatics analysis. The functional enrichment, pathway analysis, and protein-protein interaction were performed for finding key gene nodes.

## Methods

### Data resources

Gene expression profile dataset GSE152075 was obtained from the GEO database (http://www.ncbi.nlm.nih.gov/geo/). The data was produced by using an Illumina NextSeq 500 (Homo sapiens) (University of Washington, Seattle, WA). Our study analyzed five positive samples (GSM4602241, GSM4602242, GSM4602243, GSM4602244, GSM4602245) and five negative control samples (GSM4602672, GSM4602673, GSM4602674, GSM4602675, GSM4602676).

### Data acquisition and preprocessing

The raw microarray data files between SARS-CoV2 positive samples and negative controls were subsequently conducted by R script. We used a classical t test to identify DEGs with P<.05 and fold change ≥1.5 as being statistically significant.

### Gene ontology (GO) and pathway enrichment analysis of DEGs

Gene ontology (GO) analysis is a widely used approach to annotate genomic data and identify characteristic biological information. The Kyoto Encyclopedia of Genes and Genomes (KEGG) database is commonly used for systematic analysis of gene functions and annotation of biological pathways. GO analysis and KEGG pathway enrichment analysis of DEGs in our study were analyzed by the Database for Annotation, Visualization, and Integrated Discovery (DAVID) (http://david.ncifcrf.gov/) online tools. P<.05 and gene counts >10 were considered statistically significant.

### Module analysis

The Molecular Complex Detection (MCODE) was used to detect densely connected regions in PPI networks. The significant modules were from the constructed PPI network using MCODE. the functional and pathway enrichment analyses were performed using Reactome Pathway Database (https://reactome.org/).

## Results

### Identification of DEGs in COVID-19 patients

To gather further insights on host responses to COVID-19 infection, the modular transcriptional signature of COVID-19 patients was compared to that of negative controls. A total of 1353 genes were identified to be differentially expressed in COVID-19 samples with the threshold of P<0.05. Among these DEGs, 21 were up-regulated and 1332 down-regulated in COVID-19 compared with negative samples. The top 10 up- and down-regulated genes for COVID-19 and negative samples are list in table 1.

**Table 1.**

| **Entrez gene** | **Gene Symble** | **Fold-change** | **Regulation** |
| --- | --- | --- | --- |
| Top 10 down-regulated DEGs in COVID-19 |  |  |  |
| 81570 | CLPB | -1.81782581 | Down |
| 27154 | BRPF3 | -1.79174488 | Down |
| 23192 | ATG4B | -1.77926994 | Down |
| 139231 | FAM199X | -1.76768792 | Down |
| 2055 | CLN8 | -1.76573554 | Down |
| 6059 | ABCE1 | -1.75879956 | Down |
| 8310 | ACOX3 | -1.74831455 | Down |
| 23015 | GOLGA8A | -1.74438682 | Down |
| 26224 | FBXL3 | -1.73104276 | Down |
| 83440 | ADPGK | -1.72935876 | Down |
| Top 10 up-regulated DEGs in COVID-19 |  |  |  |
| 2359 | FPR3 | 1.60611739 | up |
| 389179 | EEF1A1P8 | 1.4217111 | up |
| 10563 | CXCL13 | 1.41421356 | up |
| 3034 | HAL | 1.38676074 | up |
| 2214 | FCGR3A | 1.35034198 | up |
| 4283 | CXCL9 | 1.33438197 | up |
| 342510 | CD300E | 1.32780311 | up |
| 2633 | GBP1 | 1.26094819 | up |
| 9586 | CREB5 | 1.2542903 | up |
| 168537 | GIMAP7 | 1.25151201 | up |

### Enrichment analysis of DEGs in COVID-19 patients

To further analyze the biological roles of the DEGs from negative controls versus COVID-19 positive samples, we performed KEGG pathway and GO categories enrichment analysis. KEGG pathways (http://www.genome.jp/kegg/) is an encyclopedia of genes and genomes, which was used to define the functional meanings to genes and genomes both at the molecular and higher levels^16^. In our study, the top enriched biological pathways associated with COVID-19 included “Biosynthesis of unsaturated fatty acids”, “Fatty acid elongation”, “ABC transporters”, “Lysosome”, “Vasopressin-regulated water reabsorption”, “RNA degradation” and “Fatty acid metabolism” (Figure 1).

**Figure 1.**
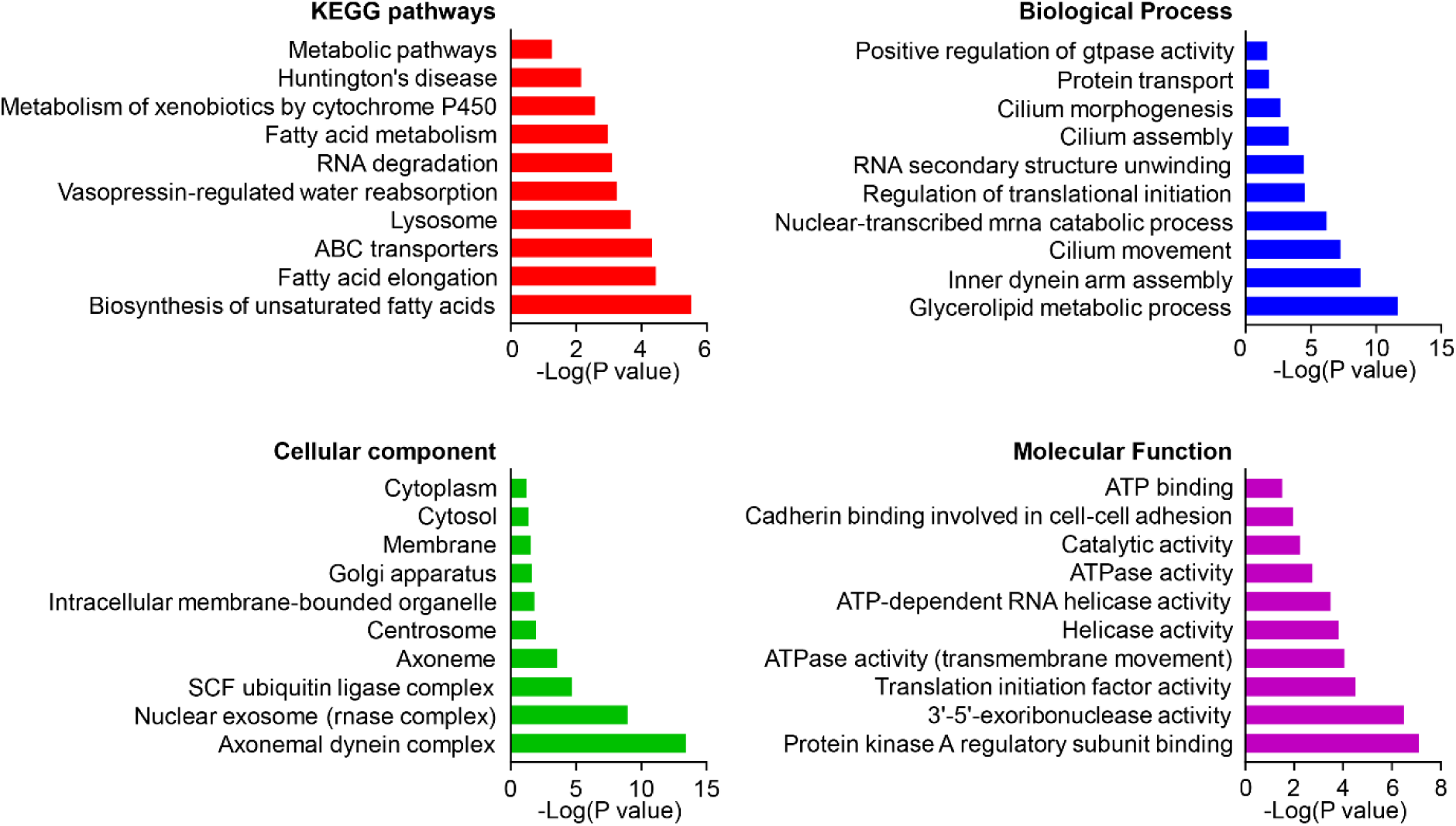
The KEGG pathways, biological process, cellular component, and molecular function terms enriched by the DEGs between COVID-19 patients and negative controls. DEGs =differentially expressed genes, KEGG = Kyoto Encyclopedia of Genes and Genomes.

Gene ontology (GO) analysis provides defined GO terms to genes, which covers cellular components, molecular functions, and biological processes^17^. We herein identified top 10 cellular components: “axonemal dynein complex”, “nuclear exosome (RNase complex)”, “SCF ubiquitin ligase complex”, “axoneme”, “centrosome”, “intracellular membrane-bounded organelle”, “Golgi apparatus”, “membrane”, “cytosol” and “cytoplasm” (Figure 1). We also identified top 10 molecular functions: “protein kinase A regulatory subunit binding”, “3’-5’-exoribonuclease activity”, “translation initiation factor activity”, “ATPase activity (transmembrane movement)”, “helicase activity”, “ATP-dependent RNA helicase activity”, “ATPase activity”, “catalytic activity”, “cadherin binding involved in cell-cell adhesion”, “ATP binding” (Figure 1). We then identified top 10 biological processes: “glycerolipid metabolic process”, “inner dynein arm assembly”, “cilium movement”, “exonucleolytic nuclear-transcribed mRNA catabolic process involved in deadenylation-dependent decay”, “regulation of translational initiation”, “RNA secondary structure unwinding”, “cilium assembly”, “cilium morphogenesis”, “protein transport” and “positive regulation of GTPase activity” (Figure 1).

### PPI network analysis of DEGs

To further explore the relationships of GGEs at the protein level, the PPI networks were constructed by using the Cytoscape software. We set the predefined criterion of combined score >0.7, a total of 2858 interactions and 1248 nodes were created to form the PPI network between negative controls and COVID-19 positive samples. Among these nodes, the top 10 hub genes with highest degree scores are shown in Table 2.

**Table 2.** Top ten genes demonstrated by connectivity degree in the PPI network

| Gene Symbol | Gene title | Degree |
| --- | --- | --- |
| ELOC | elongin C | 49 |
| DYNLL1 | dynein light chain LC8-type 1 | 48 |
| FBXL3 | F-box and leucine rich repeat protein 3 | 42 |
| FBXW11 | F-box and WD repeat domain containing 11 | 40 |
| FBXO27 | F-box protein 27 | 38 |
| FBXO44 | F-box protein 44 | 38 |
| FBXO32 | F-box protein 32 | 38 |
| FBXO31 | F-box protein 31 | 38 |
| FBXO9 | F-box protein 9 | 38 |
| CUL2 | cullin 2 | 38 |

### Module analysis

The top two significant modules of COVID-19 versus control samples were selected to analyze the functional annotation of the genes (Figure 2). We then analyzed the functional genes by using Reactome Pathway Database (https://reactome.org/). We identified top 10 significant pathways including “Anchoring of the basal body to the plasma membrane”, “Cilium Assembly”, “ Organelle biogenesis and maintenance”, “Loss of Nlp from mitotic centrosomes”, “Loss of proteins required for interphase microtubule organization from the centrosome”, “AURKA Activation by TPX2”, “Recruitment of mitotic centrosome proteins and complexes”, “Centrosome maturation”, “Regulation of PLK1 Activity at G2/M Transition” and “Recruitment of NuMA to mitotic centrosomes” in Supplemental table S1.

**Figure 2.**
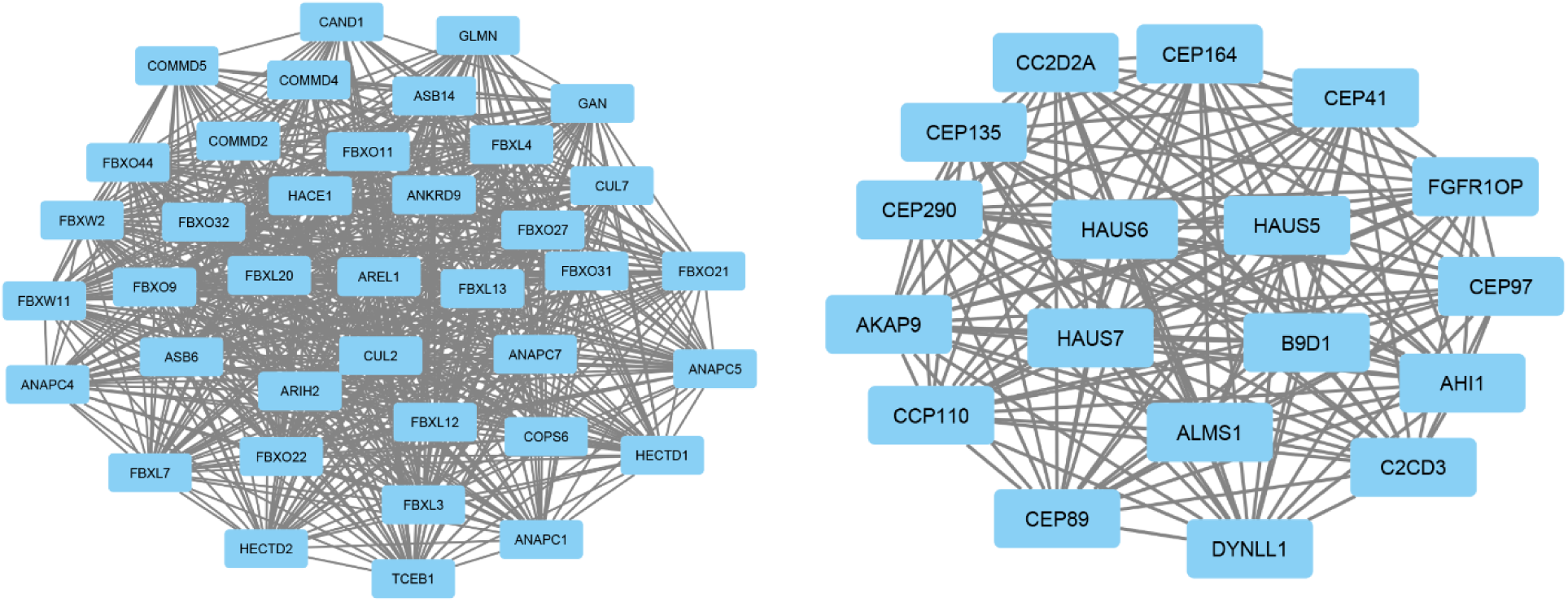
Top 2 modules from the protein-protein interaction network between COVID-19 patients and negative controls.

## Discussion

The COVID-19 has had profound economic and public health impact globally^18^. Understanding the pathogenesis of COVID-19 is critical for diagnosis and drug development. The high-throughput sequencing provides thousands of gene information which can be used to predict the potential therapeutic targets^19^.

COVID-19 has great impact on metabolic syndrome according to our analysis of KEGG pathways. The results showed six in ten most enriched pathways are involved in the regulation of fatty-related metabolism such as “Biosynthesis of unsaturated fatty acids”, “fatty acid elongation”, “ABC transporters”, “Fatty acid metabolism”, “Metabolism of xenobiotics by cytochrome P450” and “Metabolic pathways”. Many patients with COVID-19 have related to metabolic syndrome. In a study conducted in the U.S., 12% of the 1482 patients with COVID-19 had history of comorbidities. Of this total, 48.3% were obese, 28.3% were diabetics and 27.8% were cardiovascular disease^20^. In this scenario, the relationship between metabolic syndrome and COVID-19 cannot be ignored. Our analysis showed COVID-19 mainly affects fatty acid synthesis, processing, and decomposition process. Similarly, Tiffany Thomas et al found the COVID-19 could alter the fatty acid metabolism consistent with altered carbon homeostasis^21^. Thus, we believed that the high mortality rate of patients with basic metabolic diseases/COVID-19 is due to that COVID-19 changed the metabolic process of fatty acids. We also believe that the simultaneous administration of drugs that regulate fatty acid metabolism during the treatment of COVID-19 may accelerate the recovery of patients.

Besides regulating metabolism such as ATPase activity, COVID-19 is involved in cilium activity. The GO analysis showed “Cilium movement”, “Cilium assembly” and “Cilium morphogenesis” play important roles in COVID-19 patients. COVID-19 causes lung infection with loss of smell, which recently has been considered as a specific symptom. Similarly, the correlation between the COVID-19 and chemosensory dysfunction was reported, suggesting that loss of smell or taste may be considered as subclinical markers^22,23^. Numerous odorant receptors locate on sensory cilia, perceive odorants, and transduce the signal to brain. Primary cilia also perceive various environmental stimuli and regulate cell functions^24,25^. Moreover, the latest pathological studies found the expression of SARS-CoV-2 antigens in the ciliated nasal epithelial cells^26,27^. Moreover, TEM studies detected the short cilia and smell restoration in patients with COVID-19. TEM studies also provide evidence that cilia can be the absorption site or entry site for viral infection^28,29^. Thus, our results further confirmed that COVID-19 could affect the cilia system to promote the entry. It is suggested that cilia regeneration therapy may be a new direction for COVID-19 infection.

The subsequent construction of the PPI network identified 10 DEGs as potential key genes involved in COVID-19. Elongin C (ELOC), which is a critical part binds to the “BC-box motif” in the VHL-box and SOCS-box protein families. The immunodeficiency virus (FIV) protein interacts with Cullin (CUL), Elongin B (ELOB), and Elongin C (ELOC) to form an E3 ubiquitination complex to induce the degradation of feline A3s^30^. This suggests that Elongin C is critical for virus binding and functions. Cytoplasmic dynein 1 (DYNLL1), acts as a motor for the intracellular retrograde motility of vesicles and organelles along microtubules, is also reported to enhance the intracellular transport of porcine circovirus type 2 (PCV2)^31^. Circadian clocks control the processes of diseases such as inflammation, aging, virus replication and the severity of infections^32–34^. FBXL3 is reported to bind to a core circadian protein cryptochrome 2 (Cry2) to promote its degradation. Thus, FBXL3 could change the circadian rhythms in COVID-19 patients. FBXW11 (F-Box And WD Repeat Domain Containing 11) is a protein associated with diseases include digital syndrome and non-specific syndromic intellectual disability^35^. Recently, a major virulence factor NSs was found to interact with FBXW11^36^. Moreover, module analysis of the PPI network of COVID-19 suggested that the development and progression of COVID-19 were related to the cilium assembly. The genes such as AHI1, AHI1, AKAP9, B9D1, C2CD3, CEP135, CEP164, CEP290, CEP41 are involved in basal body and cilium construction after analysis by String.

In conclusion, FLOC, DYNLL1, FBXL3 and FBXW11 may be the key genes for COVID-19. Furthermore, the results suggested that several biological pathways such as “fatty-related metabolism”, “Cilium movement”, “Cilium assembly” and “Cilium morphogenesis” are commonly involved in the development of COVID-19. This study thus provides further insights into the underlying pathogenesis of COVID-19, which may facilitate the diagnosis and drug development.

## Reference

1. Salas, R.N., Shultz, J.M. & Solomon, C.G. The Climate Crisis and Covid-19 - A Major Threat to the Pandemic Response. N. Engl. J. Med. (2020).

2. Braciale, T.J. & Hahn, Y.S. Immunity to viruses. Immunol. Rev. 255, 5–12 (2013).

3. Shin, M.D., et al. COVID-19 vaccine development and a potential nanomaterial path forward. Nat Nanotechnol 15, 646–655 (2020).

4. Lauer, S.A., et al. The Incubation Period of Coronavirus Disease 2019 (COVID-19) From Publicly Reported Confirmed Cases: Estimation and Application. Ann. Intern. Med. 172, 577–582 (2020).

5. Sempowski, G.D., Saunders, K.O., Acharya, P., Wiehe, K.J. & Haynes, B.F. Pandemic Preparedness: Developing Vaccines and Therapeutic Antibodies ForCOVID-19. Cell 181, 1458–1463 (2020).

6. Giansanti, D. The Italian Fight Against the COVID-19 Pandemic in the Second Phase: The Renewed Opportunity of Telemedicine. Telemed. J. E Health (2020).

7. Robson, B. COVID-19 Coronavirus spike protein analysis for synthetic vaccines, a peptidomimetic antagonist, and therapeutic drugs, and analysis of a proposed achilles’ heel conserved region to minimize probability of escape mutations and drug resistance. Comput. Biol. Med. 121, 103749 (2020).

8. Schoeman, D. & Fielding, B.C. Coronavirus envelope protein: current knowledge. Virol. J. 16, 69 (2019).

9. Schlake, T., Thess, A., Fotin-Mleczek, M. & Kallen, K.J. Developing mRNA-vaccine technologies. RNA Biol. 9, 1319–1330 (2012).

10. DeFilippis, V., Raggo, C., Moses, A. & Fruh, K. Functional genomics in virology and antiviral drug discovery. Trends Biotechnol. 21, 452–457 (2003).

11. Wong, J.P., et al. Current and future developments in the treatment of virus-induced hypercytokinemia. Future Med Chem 9, 169–178 (2017).

12. Yang, P. & Wang, X. COVID-19: a new challenge for human beings. Cell. Mol. Immunol. 17, 555–557 (2020).

13. Tarca, A.L., Romero, R. & Draghici, S. Analysis of microarray experiments of gene expression profiling. Am. J. Obstet. Gynecol. 195, 373–388 (2006).

14. Carter, L.J., et al. Assay Techniques and Test Development for COVID-19 Diagnosis. ACS Cent Sci 6, 591–605 (2020).

15. Ward, K. Microarray technology in obstetrics and gynecology: a guide for clinicians. Am. J. Obstet. Gynecol. 195, 364–372 (2006).

16. Kanehisa, M. & Goto, S. KEGG: kyoto encyclopedia of genes and genomes. Nucleic Acids Res. 28, 27–30 (2000).

17. Hill, D.P., Smith, B., McAndrews-Hill, M.S., & Blake, J.A. Gene Ontology annotations: what they mean and where they come from. BMC Bioinformatics 9 Suppl 5, S2 (2008).

18. Shokri, T., et al. Facial Plastic and Reconstructive Surgery During the COVID-19 Pandemic: Implications in Craniomaxillofacial Trauma and Head and Neck Reconstruction. Ann. Plast. Surg. 85, S166–S170 (2020).

19. Wu, C., et al. Analysis of therapeutic targets for SARS-CoV-2 and discovery of potential drugs by computational methods. Acta Pharm Sin B (2020).

20. Garg, S., et al. Hospitalization Rates and Characteristics of Patients Hospitalized with Laboratory-Confirmed Coronavirus Disease 2019 - COVID-NET, 14 States, March 1-30, 2020. MMWR Morb. Mortal. Wkly. Rep. 69, 458–464 (2020).

21. Thomas, T., et al. COVID-19 infection alters kynurenine and fatty acid metabolism, correlating with IL-6 levels and renal status. JCI Insight 5(2020).

22. Giacomelli, A., et al. Self-reported Olfactory and Taste Disorders in Patients With Severe Acute Respiratory Coronavirus 2 Infection: A Cross-sectional Study. Clin. Infect. Dis. 71, 889–890 (2020).

23. Li, W., Li, M. & Ou, G. COVID-19, cilia, and smell. FEBS J. (2020).

24. Glezer, I. & Malnic, B. Olfactory receptor function. Handb Clin Neurol 164, 67–78 (2019).

25. Reiter, J.F. & Leroux, M.R. Genes and molecular pathways underpinning ciliopathies. Nat. Rev. Mol. Cell Biol. 18, 533–547 (2017).

26. Lukassen, S., et al. SARS-CoV-2 receptor ACE2 and TMPRSS2 are primarily expressed in bronchial transient secretory cells. EMBO J. 39, e105114 (2020).

27. Sungnak, W., et al. SARS-CoV-2 entry factors are highly expressed in nasal epithelial cells together with innate immune genes. Nat. Med. 26, 681–687 (2020).

28. Afzelius, B.A. Ultrastructure of human nasal epithelium during an episode of coronavirus infection. Virchows Arch. 424, 295–300 (1994).

29. Rautiainen, M., Nuutinen, J., Kiukaanniemi, H. & Collan, Y. Ultrastructural changes in human nasal cilia caused by the common cold and recovery of ciliated epithelium. Ann. Otol. Rhinol. Laryngol. 101, 982–987 (1992).

30. Gu, Q., et al. Identification of a Conserved Interface of Human Immunodeficiency Virus Type 1 and Feline Immunodeficiency Virus Vifs with Cullin 5. J. Virol. 92(2018).

31. Theerawatanasirikul, S., Phecharat, N., Prawettongsopon, C., Chaicumpa, W. & Lekcharoensuk, P. Dynein light chain DYNLL1 subunit facilitates porcine circovirus type 2 intracellular transports along microtubules. Arch. Virol. 162, 677–686 (2017).

32. Yuan, G., et al. Clock mutant promotes osteoarthritis by inhibiting the acetylation of NFkappaB. Osteoarthritis Cartilage 27, 922–931 (2019).

33. Yuan, G., et al. Clock mediates liver senescence by controlling ER stress. Aging 9, 2647–2665 (2017).

34. Ray, S. & Reddy, A.B. COVID-19 management in light of the circadian clock. Nat. Rev. Mol. Cell Biol. (2020).

35. Holt, R.J., et al. De Novo Missense Variants in FBXW11 Cause Diverse Developmental Phenotypes Including Brain, Eye, and Digit Anomalies. Am. J. Hum. Genet. 105, 640–657 (2019).

36. Kainulainen, M., Lau, S., Samuel, C.E., Hornung, V. & Weber, F. NSs Virulence Factor of Rift Valley Fever Virus Engages the F-Box Proteins FBXW11 and beta-TRCP1 To Degrade the Antiviral Protein Kinase PKR. J. Virol. 90, 6140–6147 (2016).

